# High molecular weight DNA extraction strategies for long-read sequencing of complex metagenomes

**DOI:** 10.1101/2021.03.03.433801

**Authors:** Florian Trigodet, Karen Lolans, Emily Fogarty, Alon Shaiber, Hilary G. Morrison, Luis Barreiro, Bana Jabri, A. Murat Eren

**Author notes:** Equal contribution (Florian Trigodet, Karen Lolans).

## Abstract

By offering extremely long contiguous characterization of individual DNA molecules, rapidly emerging long-read sequencing strategies offer comprehensive insights into the organization of genetic information in genomes and metagenomes. However, successful long-read sequencing experiments demand high concentrations of highly purified DNA of high molecular weight (HMW), which limits the utility of established DNA extraction kits designed for short-read sequencing. Challenges associated with input DNA quality intensify further when working with complex environmental samples of low microbial biomass, which requires new protocols that are tailored to study metagenomes with long-read sequencing. Here, we use human tongue scrapings to benchmark six HMW DNA extraction strategies that are based on commercially available kits, phenol-chloroform (PC) extraction, and agarose encasement followed by agarase digestion. A typical end goal of HMW DNA extractions is to obtain the longest possible reads during sequencing, which is often achieved by PC extractions as demonstrated in sequencing of cultured cells. Yet our analyses that consider overall read-size distribution, assembly performance, and the number of circularized elements found in sequencing results suggest that non-PC methods may be more appropriate for long-read sequencing of metagenomes.

## Introduction

High-throughput sequencing of metagenomes offers unprecedented insights into the diversity and gene pool of naturally occurring microbes and viruses that occupy soils (Nesme et al. 2016), marine habitats (Sunagawa et al. 2015; Gregory, Zayed, et al. 2019) and host-associated environments (Human Microbiome Project Consortium 2012; Gregory, Zablocki, et al. 2019). The high accuracy and the high throughput of the modern sequencing platforms are afforded by read lengths that typically remain below 250 bases. These relatively short reads pose significant constraints on the data utility, especially in metagenomics (Wommack, Bhavsar, and Ravel 2008).

By stitching together the short reads that partially overlap, metagenomic assembly can reconstruct orders of magnitude longer contiguous segments of input DNA (Nurk et al. 2017) and enable the recovery of microbial genomes from metagenomes (Tyson et al. 2004). In recent years, this strategy has become a primary tool in microbiology to study the ecology and evolution of naturally occurring microbial populations (Spang et al. 2015; Hug et al. 2016; Delmont et al. 2018; Pasolli et al. 2019; Al-Shayeb et al. 2020). But metagenomic assembly is inherently a challenging task (Ayling, Clark, and Leggett 2020) and the assembly of complex environments often leads to highly fragmented assemblies (Olson et al. 2019). These fragmented assemblies increase the likelihood of generating composite genomes that include contigs from multiple distinct populations (Chen et al. 2020), which risk erroneous insights into microbial ecology and evolution (Shaiber and Eren 2019; Chen et al. 2020).

By circumventing the problems associated with short read assembly, long-read sequencing offers a compelling solution to the ideal of reconstructing complete genomes from metagenomes (Driscoll et al. 2017; White et al. 2016). Nanopore sequencing (Wang, Yang, and Wang 2014) in particular has gained popularity among researchers as this strategy (1) poses no theoretical limit on read length, (2) provides new analytical opportunities with ultra-long sequences (Rand et al. 2017; Simpson et al. 2017), and (3) is accessible through affordable and easy to operate sequencing devices, such as MinION by Oxford Nanopore Technologies (“The Long View on Sequencing” 2018). Despite the high error rates and relatively lower sequencing depth, long reads from nanopore sequencing of metagenomes led to key insights from challenging systems (Reveillaud et al. 2019; Pessi et al. 2020) and enabled the recovery of circular, complete genomes from metagenomes (Somerville et al. 2018; Sanderson et al. 2018; Nicholls et al. 2019; Moss, Maghini, and Bhatt 2020; Cusco et al. 2020; Singleton et al. 2021).

The efficacy of long-read sequencing heavily depends on the structural integrity of the input DNA (Schalamun et al. 2019), which poses a new and significant challenge. Commercial DNA extraction kits that emerged during the era of short-read sequencing typically include steps that physically disrupt cells through mechanical lysis and generate highly fragmented DNA molecules. While these comercial kits improve short-read sequencing outcomes as they ensure maximum yield and coverage of DNA in a sample, they set a critical limit to the outcomes of long-read sequencing. Hence, establishing DNA extraction strategies that afford (1) preservation of high molecular weight (HMW) molecules, (2) high degree of sample purity, and (3) increased overall DNA yields has become a critical consideration for successful applications of long-read sequencing.

Phenol-chloroform DNA extractions, first popularized back in 1989 (Sambrook, Fritsch, and Maniatis 1989), have been making a resurgence as a ‘go-to’ method for extracting HMW DNA (Maghini et al. 2020). While recent studies have used this approach to recover ultra-long DNA fragments (*e.g.*, >100 kbp) from cultured organisms (Kinoshita et al. 2020; Cicha et al. 2020; Hosoe et al. 2020; Takeshita, Jang, and Kikuchi 2020; Tippelt et al. 2020), the utility of phenol-chloroform extractions for metagenomics is not yet clear. In parallel, long-read sequencing surveys of metagenomes have largely focused on high microbial biomass samples including human stool (Moss, Maghini, and Bhatt 2020) or activated sludge (Singleton et al. 2021), and best practices to study metagenomes of lower biomass samples are yet to emerge. Increasing the breadth of long-read sequencing requires DNA extraction protocols that can both produce long reads and can scale to a range of systems, including those that are associated with low microbial biomass.

Here we designed six HMW DNA extraction protocols and used an Oxford Nanopore MinION sequencer to benchmark DNA yield, host contamination, fragment size distribution, microbial taxonomy, and metagenomic assembly outcomes we obtained from each extraction method and multiple sequencing runs. We chose the human oral cavity as a model system, since it offers a challenging environment to benchmark DNA extraction strategies as it is home to complex microbial communities (Dewhirst et al. 2010) with relatively low biomass (Duran-Pinedo and Frias-Lopez 2015) and typically mixed with eukaryotic host DNA (Marotz et al. 2018).

## Results

The DNA extraction protocols we benchmarked here include four protocols based on three commercially available Qiagen DNA extraction kits (each incorporated modifications to their cell lysis procedures): (M1) the Qiagen DNeasy PowerSoil with a modified bead beating step or (M2) with bead beating replaced by enzymatic cell lysis; (M3) the Qiagen DNeasy UltraClean Microbial Kit using the manufacturer’s alternative lysis procedure to reduce DNA shearing; and (M4) the Qiagen Genomic Tip 20/G extraction kit augmented with additional enzymatic cell lysis. The remaining two protocols included in our study are (M5) a phenol-chloroform protocol, and (M6) a pulsed-field gel electrophoresis (PFGE)-based agarose encasement extraction protocol followed by agarase digestion. Throughout the text we refer to these protocols as M1, M2, M3, M4, M5 and M6, and we denote technical replicates as “MX_1” or “MX_2”. To benchmark these protocols that are detailed in the Methods section, we used a tongue dorsum sample pooled together from 13 samples collected from the same individual.

### Yield and quality metrics of isolated DNA vary between protocols

To ascertain the suitability for long-read sequence analysis, we first analyzed the quantity and quality of DNA isolated from each extraction protocol using both fluorometric (Qubit) and spectrophotometric (Nanodrop) methods (Table 1). The fluorescent dye used by Qubit binds specifically to its target molecule, dsDNA, and provides the most accurate DNA quantitation. DNA concentrations were comparable between technical replicates for all methods, and three extraction protocols (M4, M5 and M6) were distinguished from the others with concentrations >75 µg/ml. M4 exhibited the highest sample concentrations with a mean of 110 µg/ml. In comparison, the mean concentrations from M1, M2 and M3 were far less at 2.03, 4.15 and 33.45 µg/ml, respectively. The MinION sequencing protocol for the Ligation Sequencing Kit (SQK-LSK108) advises using 1 – 1.5 µg DNA. As we resuspended DNA into a final volume of 100 µl, only M3 to M6 had sufficient DNA yield to meet the recommended input.

**Table 1.**
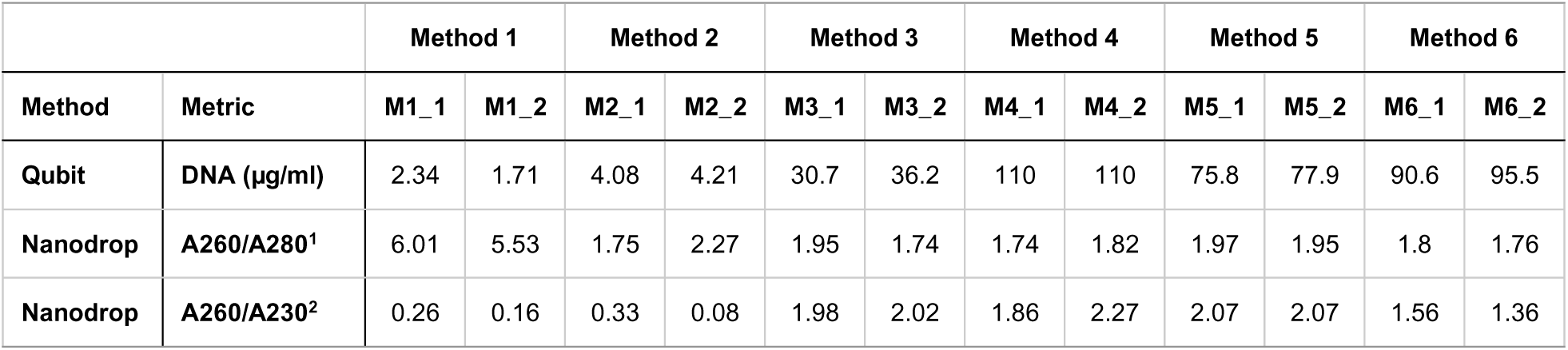
Summary of DNA concentration and quality metrics. Technical replicates are denoted as “MX_1” and “MX_2”. 1 Primary measure of nucleic acid purity –Expected value for “pure” DNA: ∼1.8. Values significantly lower are not reliable and >2 indicates RNA contamination. 2 Secondary measure of nucleic acid purity - evaluates residual chemical contamination (phenol, guanidine HCL, carbohydrate carryover). Expected values are in the range of 1.8 – 2.2.

We then considered the absorption spectra to assess sample purity and identify potential non-nucleic acid contamination. The A260/A280 ratio indicates DNA purity with expected values around 1.8 for pure DNA. All extraction methods fell within the desired range (1.74 - 1.97), except for both M1 replicates and one replicate of M2; however, samples with concentrations approaching the lower limit of 2 µg/mL may result in unacceptable 260/280 ratios. The A260/A230 ratio signifies possible residual chemical contamination such as EDTA, phenol, guanidine salts (often used in column-based kits), or carbohydrate carryover. M3, M4 and M5 had ratios close to the expected value of 2. Overall, M4 had the best combined yield and purity metrics; M5 and M6 were deemed suitable alternatives. The purity metrics and congruence of Qubit and Nanodrop concentrations in M3 were also desirable, yet ∼2-3-fold lower DNA yield proved M3 less appealing.

We ran an agarose gel electrophoresis to visually assess the crude DNA fragment size distribution (Figure 1). All DNA samples migrated predominantly as a single high-molecular-weight DNA band that aligned with (or was larger) than the reference 23.1 kbp fragment of HindIII-digested lambda DNA. A light smear, visible to 2.0 kb, was observed in M4 indicating the presence of smaller fragments. Recovery of larger DNA fragments by M5 was denoted by the predominant band running higher than the 23.1 kbp fragment. Low DNA yield can result in sample loss during library preparation or low pore occupancy on MinION during sequencing. While working with samples of low DNA yield is inevitable, spiking in known DNA (such as lambda DNA) to ‘pad’ samples with low DNA yields may be used to start sequencing as we demonstrated before (Reveillaud et al. 2019). But to minimize the need for additional DNA to ‘pad’ samples with low DNA yield due to the extraction protocol, we eliminated protocols M1, M2 and M3 from any further evaluation as they consistently resulted in mediocre DNA yield.

**Figure 1.**
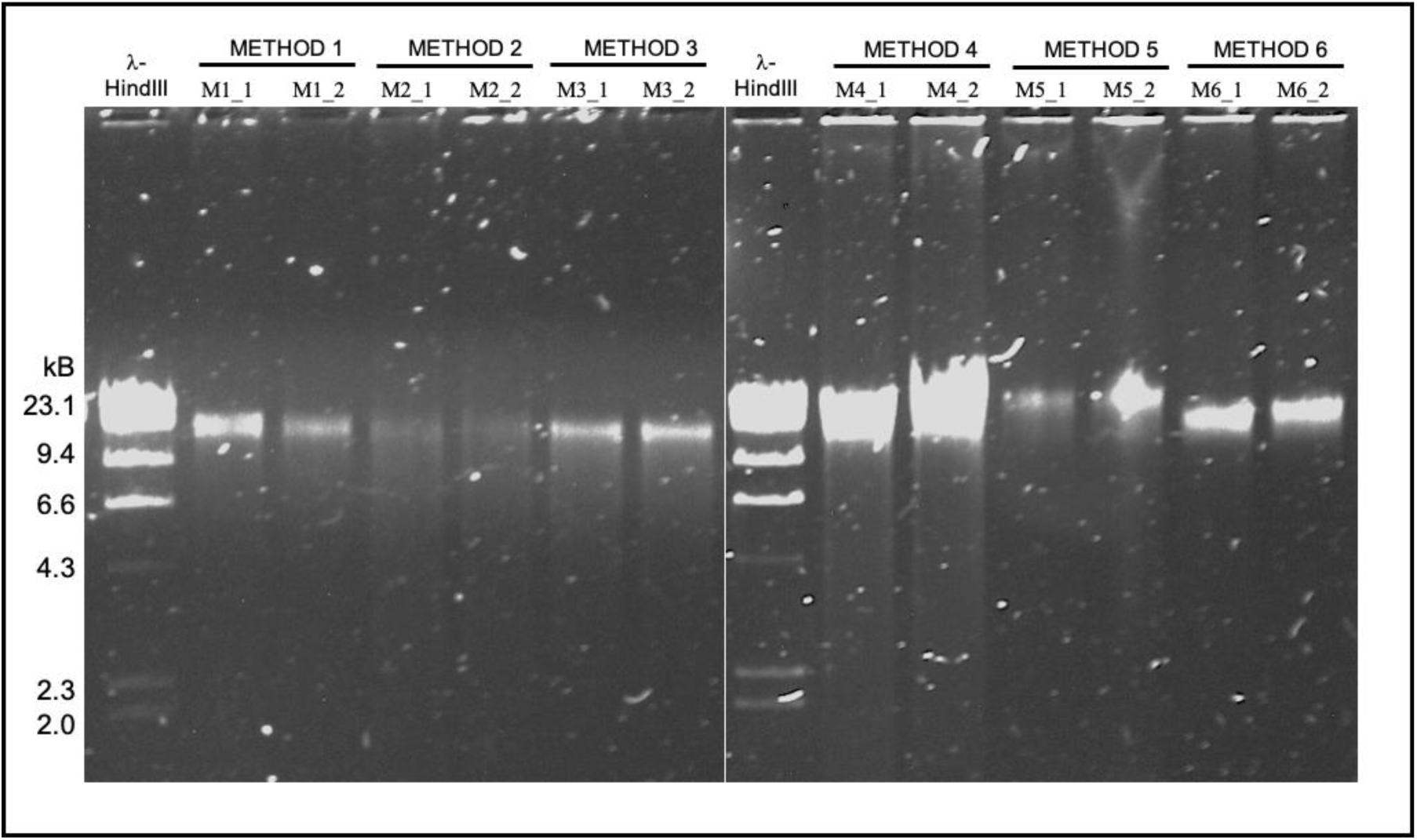
Agarose gel electrophoresis of genomic DNA isolated from a pool of tongue dorsum samples. Genomic DNA was electrophoresed on a 0.8% (w/v) agarose gel. Method 1, Method 2 and Method 3 replicates (22 ng input, left panel) and Method 4, Method 5 and Method 6 replicates (44 ng input, right panel) are shown. Different DNA inputs were used based on overall sample availability. λ-HindIII, Lambda DNA, digested with the restriction endonuclease HindIII, was used to assess fragment size distribution.

Next, we sequenced the DNA from M4, M5, and M6 using two MinION flow cells to ensure the replicates of the same protocol were sequenced on different runs (Run 1: M4_1, M5_1, and M6_1; Run 2: M4_2, M5_2, and M6_2). The sequencing runs generated 4.84 and 7.79 Gb, respectively. The increase in the output in the second run could be attributed to less sample loss during library preparation and subsequent input of 3X more DNA (142.8 ng versus 466.2 ng) into the flow cell. After performing a quality filtering step (using a minimum Q-score of 7), the percentage of reads passing the quality check (Pass_Reads) was similar between extraction methods within each flow cell (96% and 93-94% respectively, Table S1.a). However, a comparison between runs demonstrated that a higher percentage of sequences were removed (Fail_Reads) in the second run (Table S1.a). The total number of nucleotides was comparable between M4 and M5 within sequencing runs (1,699,213,259 and 1,641,389,967 on average, respectively), and was much smaller for M6 (949,615,901 on average) in both runs.

### Eukaryotic contamination is enriched in the pool of shorter fragments

As samples from the human oral cavity prepared for metagenomic sequencing are typically associated with extensive eukaryotic contamination which can account for up to 45% of the short-read sequencing product (Shaiber et al. 2020), we assessed the amount of host contamination in each DNA sample by mapping reads to a reference human genome from the NCBI (GRCh38). Host DNA contamination was high for all protocols. M6 had the least amount of human DNA (on average, 63%) compared to M4 (on average, 75%) and M5 (on average, 81%) (Table 2). This trend persisted when comparing their cumulative sequence lengths, although our analysis of read length distribution showed that the human reads were predominantly composed of shorter fragments (Figure 2, insets). The increased representation of human DNA in M5 compared to other extraction methods is likely due to the use of detergents versus enzymes. Lytic detergents exert their effect on both bacterial and eukaryotic cells, while lytic enzymes target bacterial cells only. M5 lacked lytic enzymes and included SDS, a stronger lytic detergent than the ones used in M4 (Tween 20, Triton X-100) and M6 (SLS, Brij, deoxycholate).

**Figure 2.**
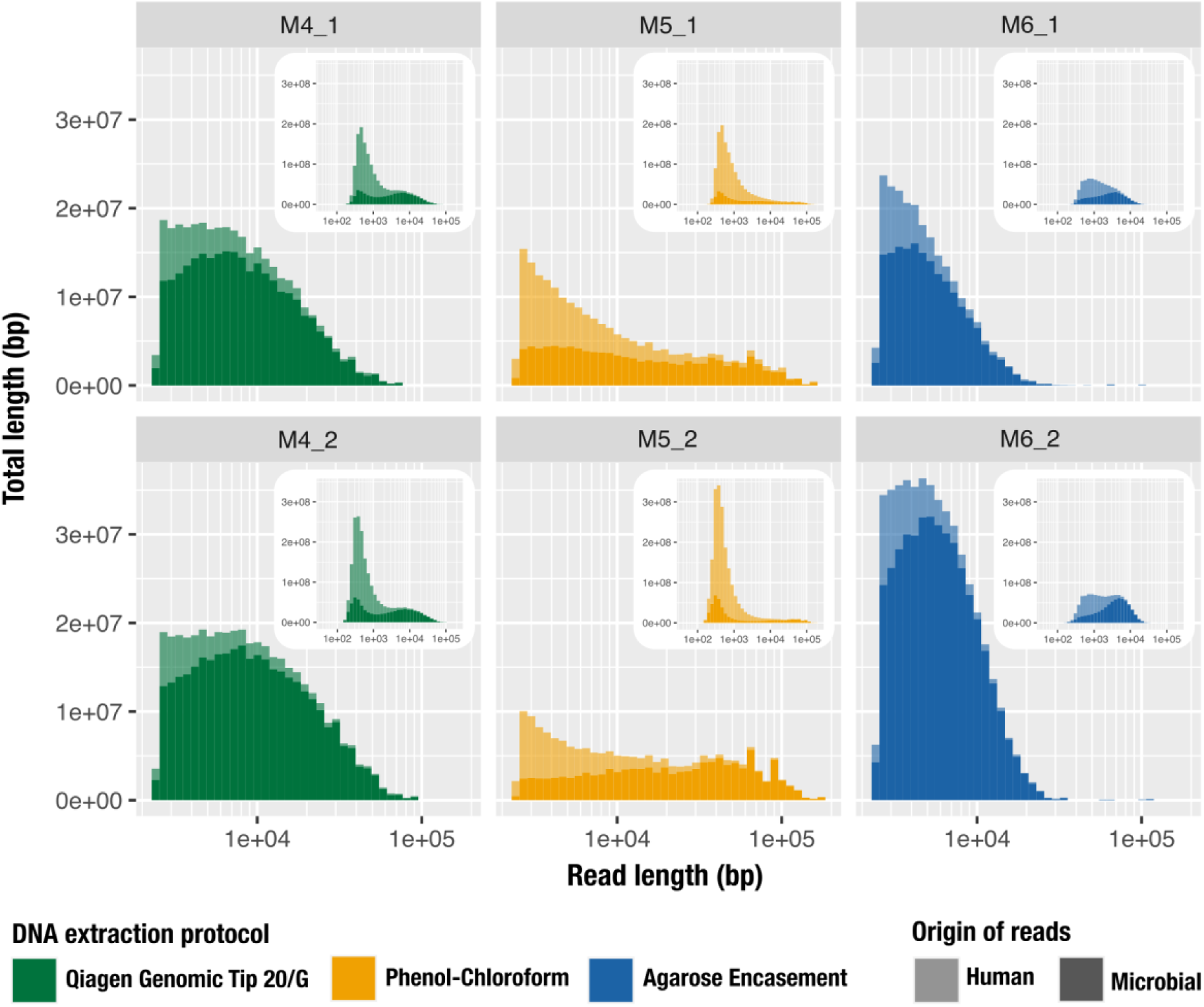
The impact of DNA extraction protocol on the distribution of human (lighter color) and microbial (darker color) read lengths from MinION sequencing. These histograms visualize the total accumulative length (total number of nucleotide sequenced) per range of individual read lengths. The x-axis represents the read length in log scale and the y-axis represents the cumulative length for a given size bin (bar width). The main panel shows the size distribution of reads >2,500 bp for M4 (green), M5 (yellow) and M6 (blue) while the inset panel shows the size distribution of all reads, using the same data. Results are outlined vertically by extraction method (replicate 1, top panel; replicate 2, lower panel).

**Table 2.** The impact of HMW DNA extraction protocol on proportional read numbers and sequence lengths according to read type (microbial versus human).

|  | METHOD 4 |  | METHOD 5 |  | METHOD 6 |  |
| --- | --- | --- | --- | --- | --- | --- |
|  | M4_1 | M4_2 | M5_1 | M5_2 | M6_1 | M6_2 |
| **All Reads** |  |  |  |  |  |  |
| Number of Reads | 2,052,300 | 3,681,083 | 2,008,795 | 4,338,575 | 739,186 | 962,065 |
| Total Length (bp) | 1,439,760,359 | 1,958,666,159 | 1,310,980,562 | 1,971,799,372 | 766,599,939 | 1,132,631,863 |
| **Human Reads** |  |  |  |  |  |  |
| Number of Reads | 1,584,229 | 2,674,013 | 1,664,447 | 3,433,184 | 490,377 | 585,733 |
| % | 77.19 | 72.64 | 82.86 | 79.13 | 66.34 | 60.88 |
| Total Length (bp) | 908,119,237 | 1,199,195,722 | 1,025,965,072 | 1,535,632,081 | 404,681,613 | 463,311,606 |
| % | 63.07 | 61.23 | 78.26 | 77.88 | 52.79 | 40.91 |
| **Microbial Reads** |  |  |  |  |  |  |
| Number of Reads | 468,071 | 1,007,070 | 344,348 | 905,391 | 248,809 | 376,332 |
| % | 22.81 | 27.36 | 17.14 | 20.87 | 33.66 | 39.12 |
| Total Length (bp) | 531,641,122 | 759,470,437 | 285,015,490 | 436,167,291 | 361,918,326 | 669,320,257 |
| % | 36.93 | 38.77 | 21.74 | 22.12 | 47.21 | 59.09 |

### Read size distribution is not uniform across extraction methods

After the removal of sequences that match the human genome, we assumed that the vast majority of the sequences were of microbial origin. We further focused on sequences that were longer than 2,500 nucleotides, and quantified the number of reads and their length distribution across M4, M5, and M6 (Table 3).

**Table 3.** Microbial read size distribution. All percentages are relative to the total reads (or total length) of the quality filtered reads, prior to removal of human reads.

|  | Method 4 |  | Method 5 |  | Method 6 |  |
| --- | --- | --- | --- | --- | --- | --- |
|  | M4_1 | M4_2 | M5_1 | M5_2 | M6_1 | M6_2 |
| **All Reads** |  |  |  |  |  |  |
| Number of Reads | 2,052,300 | 3,681,083 | 2,008,795 | 4,338,575 | 739,186 | 962,065 |
| Total Length (bp) | 1,439,760,359 | 1,958,666,159 | 1,310,980,562 | 1,971,799,372 | 766,599,939 | 1,132,631,863 |
| **All Microbial Reads** |  |  |  |  |  |  |
| Number of Reads | 468,071 | 1,007,070 | 344,348 | 905,391 | 248,809 | 376,332 |
| % | 22.81 | 27.36 | 17.14 | 20.87 | 33.66 | 39.12 |
| Total Length (bp) | 531,641,122 | 759,470,437 | 285,015,490 | 436,167,291 | 361,918,326 | 669,320,257 |
| % | 36.93 | 38.77 | 21.74 | 22.12 | 47.21 | 59.09 |
| N50 | 2,810 | 1,929 | 1,116 | 449 | 2,524 | 3,649 |
| L50 | 38,513 | 62,126 | 34,874 | 155,142 | 38,768 | 52,182 |
| Longest Microbial Reads (bp) | 73,029 | 92,515 | 163,320 | 180,460 | 68,189 | 116,730 |
| **Microbial Reads > 2.5 kb** |  |  |  |  |  |  |
| Number of Reads | 43,173 | 49,572 | 13,752 | 10,182 | 39,270 | 82,221 |
| % | 2.10 | 1.35 | 0.68 | 0.23 | 5.31 | 8.55 |
| Total Length (bp) | 278,184,355 | 352,174,239 | 108,718,450 | 116,813,051 | 182,222,365 | 425,589,257 |
| % | 19.32 | 17.98 | 8.29 | 5.92 | 23.77 | 37.58 |

Our comparison of the five longest reads per replicate revealed that M5 produced the longest fragments (130,355 to 180,460 bp) while M4 ranged from 68,275 to 92,515 bp and M6 had a single 116,730 bp read followed by significantly shorter reads (28,218 to 68,189 bp) (Table 3). The replicates of M5 yielded 24 and 38 reads that were over 100,000 bp (Table S1.b), but this extraction method was also associated with the smallest N50 score due to the large fraction of shorter reads (Figure 2, insets). The contribution of reads above 2,500 bp to the total sequencing length (total number of nucleotides) was greater in M4 (mean, 18.65%) and M6 (mean, 30.68%), while dropping to 7.11% for M5 (Table 3, Figure 2). M6 had a few short reads contributing to the total sequencing length, but also lacked long reads as there were only 40 and 186 reads above 20 kbp in the two runs (Table S1.b). We find it surprising that M6, an extraction strategy that can yield over a million base pair DNA fragments (Anand 1986), have produced only four reads that were longer than 50 kbp when we used identical DNA mass inputs for all extractions. We speculate that the very long fragments M6 may have produced may have been lost during library preparation steps or become stuck in the pores during sequencing.

### Size selection has limited utility and leads to substantial loss of biomass

Despite adding the nanopore recommended 1X volume of solid-phase reversible immobilization (SPRI) beads to reaction mixtures to remove small fragments during library preparation steps, there was a large contribution of shorter reads (< 2500 bp) in our initial sequencing effort. Thus, we evaluated additional options to reduce their numbers and increase the representation of longer fragments. We sought to determine the effect of using the BluePippin system on fragment size metrics by utilizing the ‘high-pass collection mode’, which enables the collection of DNA molecules above a certain user-defined size. To maximize sequences that might contain ribosomal RNA genes, which are particularly useful for taxonomic assignments (Camanocha and Dewhirst 2014), we chose 6 kbp as our minimum threshold value. Due to its ample material availability, we used replicates of M4 to compare the size-selected sequencing metrics with data from the previous untreated sequencing runs. High sample loss is a known drawback of BluePippin high-pass size filtering as the manufacturer warns to expect a loss between 20 - 50% of the sample input. However, we were able to recover only 16% to 18% of the 3,300 ng input DNA for each replicate after the size selection step, in agreement with low recovery rates (25% to 35%) also reported by others (Schalamun et al. 2019). Even though we started the size selection step with more than 3X the amount of DNA than was used for the untreated workflow, our recovery post-size selection was 550-600ng, meaning that our sample input into the start of library preparation was half of the DNA input that went into the untreated sequencing run. Strikingly, the number of reads and total number nucleotides was reduced by 80-90% compared to the untreated sample (Table 4, Table S1.c), a likely consequence of the reduced sample input. Other notable shifts included a reduced proportion of human contamination and an increased N50 (Table 4). The number of microbial reads above 2.5 kbp and their cumulative length were comparable for both approaches, which is quite impressive given the low amount of input DNA in the size-selected samples. However, this parallel did not persist when evaluating longer DNA fragments as the cumulative length of microbial reads above 20 kbp was greater for non-size selected samples. Propelled by vastly reduced read numbers (and despite the superior N50), size selection step did not result in the substantial improvements we hypothesized in overall read lengths and the cumulative nucleotide sequences. Considering the additional demands on (1) sample requirements, (2) reagent/personnel costs, and (3) sample handling and processing times, we consider size selection of this type to have limited utility in this context.

**Table 4.** Comparison of sequencing run read metrics between untreated and BluePippin size-selected samples. All percentages are relative to the total reads (or total length) of the quality filtered reads.

|  | Untreated |  | BluePippin High-Pass Size Selection |  |
| --- | --- | --- | --- | --- |
|  | M4_1 | M4_2 | M4_1 SS | M4_2 SS |
| **All reads** |  |  |  |  |
| Total reads | 2,052,300 | 3,681,083 | 221,344 | 430,986 |
| Total length (bp) | 1,439,760,359 | 1,958,666,159 | 409,870,430 | 450,341,701 |
| **Human reads** |  |  |  |  |
| Number of Reads | 1,584,229 | 2,674,013 | 117,071 | 282,113 |
| % | 77.19 | 72.64 | 52.89 | 65.46 |
| Total Length (bp) | 908,119,237 | 1,199,195,722 | 103,754,814 | 161,759,870 |
| % | 63.07 | 61.23 | 25.31 | 35.92 |
| **Microbial reads** |  |  |  |  |
| Number of Reads | 468,071 | 1,007,070 | 104,273 | 148,873 |
| % | 22.81 | 27.36 | 47.11 | 34.54 |
| Total Length (bp) | 531,641,122 | 759,470,437 | 306,115,616 | 288,581,831 |
| % | 36.93 | 38.77 | 74.69 | 64.08 |
| N50 | 2,810 | 1,929 | 6,106 | 5,594 |
| **Microbial Reads > 2.5 kb** |  |  |  |  |
| Number of Reads | 43,173 | 49,572 | 40,248 | 34,949 |
| % | 2.10 | 1.35 | 18.18 | 8.11 |
| Total Length (bp) | 278,184,355 | 352,174,239 | 251,096,073 | 215,008,868 |
| % | 19.32 | 17.98 | 61.26 | 47.74 |
| **Microbial Reads > 20 kb** |  |  |  |  |
| Number of Reads | 1,269 | 2,294 | 300 | 407 |
| % | 0.06 | 0.06 | 0.14 | 0.09 |
| Total Length (bp) | 34,892,531 | 66,727,907 | 7,643,616 | 11,205,696 |
| % | 2.42 | 3.41 | 1.86 | 2.49 |

### Extraction method and read length distribution alter taxonomic profiles

Extraction methods may have distinctive biases which can impact the determination of microbial community from metagenomic data, like a differential ability to lyse gram-negative versus gram-positive organisms. To investigate the microbial community composition of our long-read metagenomes we used bacterial, archaeal and eukaryotic single-copy core genes (SCGs) and ribosomal RNAs (rRNAs). Given the high rate of insertion or deletion errors in nanopore sequencing, predicting genes accurately is a significant challenge in uncorrected reads. While we assumed that these biases would similarly impact all extraction methods and elected to not correct our raw reads, we suggest the use of appropriate bioinformatics tools to correct frame-shift errors in long-read sequencing results (Arumugam et al. 2019; Huang, Liu, and Shih 2020).

In agreement with the total length of bacterial reads across extraction strategies, M4 had the most and M5 had the least number of genes predicted (Table 5). M5 also yielded the lowest number of 16S rRNA genes with an average of 123 genes as compared to M4 and M6, which yielded an average of 418 and 398 genes, respectively. Read size distribution played an important role in the recovery of SCGs and rRNAs genes: for example, M5_2 and M6_1 had a comparable total length of microbial reads (436 and 361 Mbp respectively, Table 3) but the numbers of SCGs and rRNAs genes were two times larger in M6_1, despite a smaller number of predicted genes. These two extractions strategies differed drastically in their read size distribution (Figure 2) with M6_1 having fewer very long reads (>50,000 bp, Table S1.b) but more medium-sized ones (>2,5 kbp, Table 3). To complement this observation, M6_2 had the most SCGs and rRNA genes out of all extractions and also the most reads above 2.5 kbp (Table 3), despite having very few long reads (only three > 50 kbp, Table S1.b) and being the second smallest sample in terms of number of reads and total length before the removal of human reads (Table 3).

**Table 5.** Results of prodigal gene calling and HMM hits for single-copy core genes (SCGs) and ribosomal RNAs (rRNAs). Analysis was performed after removal of human-reads.

|  | M4_1 | M4_2 | M5_1 | M5_2 | M6_1 | M6_2 |
| --- | --- | --- | --- | --- | --- | --- |
| **Number of Genes** | 676,577 | 994,050 | 391,675 | 665,986 | 440,845 | 771,653 |
| **Bacterial SCGs** | 2,893 | 3,131 | 1,309 | 1,210 | 2,136 | 3,559 |
| **Ribosomal RNAs**<br>(per 1,000 genes) | 901 (1.33) | 1,107 (1.11) | 295 (0.75) | 307 (0.46) | 655 (1.49) | 1,355 (1.76) |
| **Bacterial 16S rRNA** | 373 | 462 | 120 | 125 | 249 | 547 |

We used the Human Oral Microbiome Database (HOMD) to assign taxonomy to the 16S rRNA genes found in our long reads. The top genera included *Prevotella*, *Rothia*, *Streptococcus*, *Veillonella*, which are commonly found in oral samples (Mark Welch et al. 2016; Zaura et al. 2009). We observed a comparable taxonomic profile between M4 and M6, with *Streptococcus*, *Veillonella*, *Prevotella* and *Actinomyces* identified as the most abundant genera (Figure 3). M5 differed with relatively more *Prevotella* and *Haemophilus* (both gram negative bacteria) and less *Streptococcus* (a gram positive bacterium).

**Figure 3.**
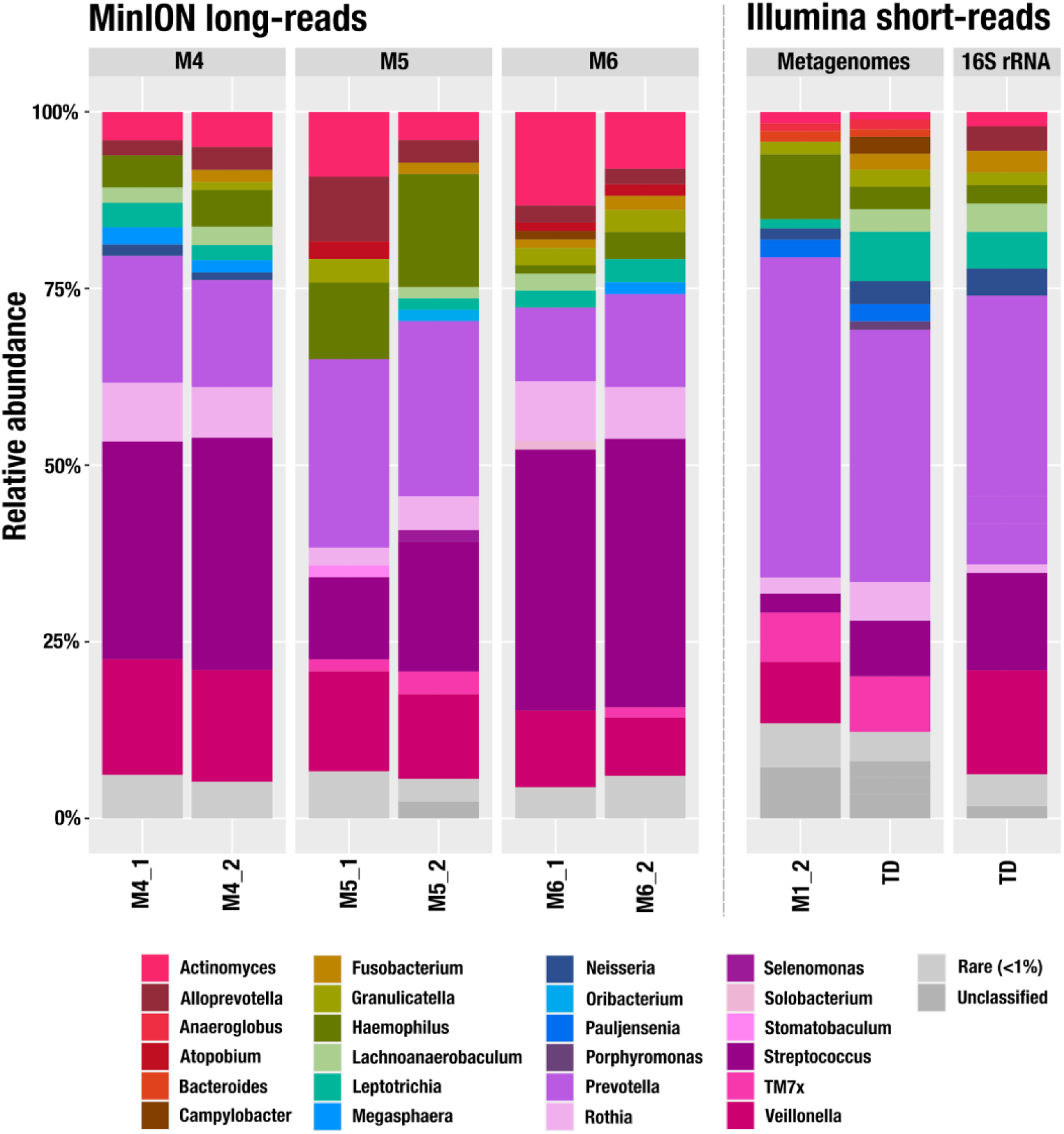
Relative abundance of 16S rRNA at the genus level. We used the Human Oral Microbiome Database (HOMD) to assign taxonomy to the 16S rRNA from the MinION reads. For the short-read metagenomes, we used the taxonomy of the ribosomal gene S7 with the Genome Taxonomy Database (GTDB). We processed the 16S rRNA amplicons with the Minimum Entropy Decomposition (MED) algorithm and used Silva v132 to assign taxonomy. Genera representing less than 1% of a sample were pooled as rare (in light grey). Samples noted as TD correspond to an additional sampling performed two weeks after the initial pool of samples used for the long-read extractions.

We then compared these profiles to the short-read sequencing of 16S rRNA gene amplicons found in a sample we collected for the same individual two weeks after the initial sampling. While long-read and amplicon sequencing results were similar to each other qualitatively, the relative abundance estimates between these approaches differed (Figure 3, Table S2). These differences can be attributed to differences in sequencing depth and the recovery of signal from the rare taxa, as well as to the different sampling time and the impact of primer biases. For instance, our 16S rRNA gene primers did not match to TM7, a prevalent taxon in the human oral cavity, which explains their absence in the amplicon sequencing.

Finally, we compared taxonomic profiles across protocols to the short-read sequencing of two metagenomes from the same individual generated from (1) the sample used for M1_2 and (2) the sample used for 16S rRNA gene amplicon sequencing (sample TD). To estimate the relative abundance of taxa in metagenomes we used short read coverage of single-copy core genes matching to ribosomal protein S7, which was the most frequently found ribosomal protein in the assemblies of short-read metagenomes (Table S3). While the genera composition was very comparable with long-read metagenomes, we observed a lower relative abundance of *Streptococcus* and higher relative abundance TM7, which is likely due to copy number of the rRNA operons in these genera (Stoddard et al. 2015) that skew the relative abundance estimates based on short-read amplicons and long-read metagenomes. We also investigated the distribution and taxonomic assignment of ribosomal proteins in long-read metagenomes (Table S3, Figure S1), however, relatively low sequencing depth of these data and challenges associated with gene calling in uncorrected MinION sequences prevented reliable insights.

Overall, these results highlight (1) the differences observed between extraction methods (*i.e.*, gram negative/positive biases), but also (2) the critical role that the read size distribution has on the recovery of SCGs, ribosomal RNAs and consequently, its impact on taxonomic profiling.

### Read size distribution dramatically influences assembly outcomes

We finally compared different extraction methods by assembling microbial reads using a long-read assembler, Flye (Kolmogorov et al. 2020). M4 and M6 resulted in larger assemblies with an average size of 31,761,136 bp and 28,578,283 bp respectively, while the average assembly size of M5 was much smaller at 11,269,288 bp (Table 6). The extraction method M4 stood out for its high N50, but also by having the longest contig assembled (1,183,656 bp) and the most contigs above 100 kbp. The number of predicted genes was directly related to the assembly size. For that reason, M4 had on average more genes predicted and more SCGs and ribosomal RNAs, which lead to more bacterial genomes expected in the assembly (based on the most frequently occurring count of SCG hits). The assembly step did not result in any circular bacterial chromosomes but there were a few circular plasmids and phage genomes. M5 had the fewest circular contigs (Table 6), and while M6 had more, they were shorter. M4 stood out again with the most circular contigs, including the largest in all assemblies (157,084 bp).

**Table 6.** Flye assembly statistics.

|  | M4_1 | M4_2 | M5_1 | M5_2 | M6_1 | M6_2 |
| --- | --- | --- | --- | --- | --- | --- |
| **Total Length (bp)** | 28,192,179 | 35,330,092 | 10,718,489 | 12,020,087 | 17,671,819 | 39,484,746 |
| **Number of Contigs** | 401 | 466 | 137 | 106 | 483 | 855 |
| Num Contigs > 5 kb | 370 | 412 | 130 | 102 | 467 | 775 |
| Num Contigs > 10 kb | 347 | 384 | 124 | 100 | 440 | 715 |
| Num Contigs > 20 kb | 299 | 336 | 108 | 97 | 356 | 570 |
| Num Contigs > 50 kb | 159 | 194 | 74 | 69 | 86 | 203 |
| Num Contigs > 100 kb | 64 | 86 | 33 | 35 | 21 | 70 |
| **Longest Contig (bp)** | 1,030,374 | 1,183,656 | 506,923 | 681,941 | 306,827 | 879,856 |
| **Shortest Contig (bp)** | 515 | 533 | 2114 | 1032 | 927 | 518 |
| **N50 (bp)** | 130,311 | 157,084 | 124,183 | 168,272 | 44,719 | 71,759 |
| **Number of Genes** | 49,565 | 61,631 | 17,062 | 19,577 | 31,650 | 69,753 |
| **Single-copy core genes** |  |  |  |  |  |  |
| Bacteria_71 | 746 | 900 | 258 | 292 | 508 | 994 |
| Archaea_76 | 393 | 496 | 123 | 166 | 305 | 568 |
| Protista_83 | 19 | 22 | 2 | 6 | 10 | 22 |
| Ribosomal RNAs | 75 | 116 | 30 | 43 | 44 | 100 |
| **Num of expected bacterial genome** | 9 | 11 | 3 | 4 | 5 | 9 |
| **Circular contigs** |  |  |  |  |  |  |
| Number of circular contigs | 20 | 32 | 8 | 5 | 9 | 24 |
| Max length | 86,674 | 157,084 | 156,592 | 157,040 | 24,661 | 88,575 |

## Discussion

Prior to undertaking this evaluation, we established two key conditions that a successful high molecular weight DNA extraction method to study host-associated environments with low microbial biomass would need to fulfill: optimizing a read length distribution profile to improve downstream sequence analyses and minimizing the proportion of host-associated DNA contamination. While the utility of minimizing non-target DNA in sequencing libraries is relatively obvious from a resource conservancy point of view, the impact of the read length distribution on studies of metagenomes may escape attention when a sequencing strategy focuses on obtaining the longest possible reads. Based on the assembly of 2,267 genomes and the average length of rDNA operons, Koren et al. had suggested 7 kbp as the ideal read length that could span through most repeats for dramatic improvements in genome assembly (Koren et al. 2013). The modified column-based extraction method (M4) consistently yielded the greatest number of reads over 10 kbp and led to most successful downstream assemblies as indicated by the total size, the length of longest contigs, and the number of circularized elements.

The representation of host DNA contamination varied between the extraction methods, and reached its maximum in the phenol-chloroform extraction (M5). As a consequence, the phenol-chloroform extraction resulted in fewer microbial reads for an equal amount of DNA input (Table S2). While host DNA was enriched in the pool of shorter sequences and thus could be eliminated through size selection, our analyses revealed a high cost for this step as size selection removed many of the very long reads (> 60 kbp) and required three times more DNA as input. A unique advantage of nanopore sequencing platforms is the real-time access to the product that is being sequenced, which promotes new methods for reference-based identification of non-target DNA molecules and their real-time rejection from an active pore to minimize their impact on sequencing results (Kovaka et al. 2020). Maximizing the input DNA quality, and the use of novel assembly algorithms (Kolmogorov et al. 2020; Koren et al. 2017; Li 2016; Ruan and Li 2020), effective polishing strategies (Loman, Quick, and Simpson 2015; Vaser et al. 2017; Walker et al. 2014; Morisse et al. 2021), will lead to new insights in metagenomics especially with complete circular MAGs.

## Conclusions

Phenol-chloroform extraction is frequently endorsed for its ability to produce extremely long DNA fragments for long-read sequencing, which was also the case in our study as M5 yielded the longest fragments we sequenced. Yet, our findings show that modified versions of commercially available column-based (M4) and agarose-based (M6) extraction methods outperform phenol-chloroform long-read sequencing of metagenomes. Our analyses suggest that the Qiagen Genomic Tip 20/G extraction kit augmented with additional enzymatic cell lysis is a promising approach to generate high-quality input DNA from complex metagenomes for long-read sequencing.

## Materials and Methods

### Tongue Dorsum Sample Collection

A single healthy individual self-collected scraping of their tongue dorsum on 13 separate occasions (1 per day) over the course of 3 weeks. We used BreathRx Gentle Tongue Scraper (Philips Sonicare) for sample collection which was performed prior to eating, drinking or performing oral hygiene. Starting as far back as possible on the tongue, the scraper was passed forward over the entire surface three sequential times. We transferred the collected material to 520 µl of PBS (phosphate-buffered saline) and immediately stored in -20°C until processing. Another self-collected tongue dorsum sample from this same individual was obtained 2 weeks later.

### DNA extraction methods

We compared six DNA extraction methods tailored for high molecular weight DNA recovery. We included both commercially available kits and non-kit methods used in the published literature which incorporated different combinations of cell lysis mechanisms and DNA purification methodologies. To facilitate direct comparison between all extraction methods, we thawed and pooled the 13 samples immediately prior to DNA extraction. After homogenizing the pooled sample by vortexing for 15-sec, we used a 500 µl aliquot as the starting material for each extraction method, which was performed in duplicate. We resuspended the isolated genomic DNA from each method in a final 100 µl volume in 1.5 ml LoBind microfuge tube (Eppendorf, Hauppauge, NY). We sought to maximize read lengths by implementing best practices for handling HMW DNA throughout all the methods. We eliminated vortexing and mixing by pipetting, when possible, in favor of end-over-end tube rotation to minimize velocity gradients. We used wide-bore pipet tips with gentle pipetting to reduce DNA breakage and avoided unnecessary freeze-thaw cycles by storing DNA at 4°C until sequence analysis.

### Method 1 (M1), DNeasy PowerSoil Isolation Kit with modified bead beating

DNeasy PowerSoil® DNA Isolation kits (Qiagen) are commonly used in metagenomics to extract high quality DNA from environmental matrices. We sought to determine its compatibility with nanopore sequencing protocols by amending the PowerSoil® DNA isolation protocol to include a modified bead beating step (Edwards et al 2019), as a way to minimize velocity gradients and reduce DNA shearing. We transferred 500 µl of the pooled sample to the kit provided PowerBead tube, which we inserted flat into IKA Works Inc MS2S8 Minishaker for Bioanalyzer DNA chips. Samples were agitated for 10 minutes (in 1 min pulse increments) at 2400 rpm (Edwards et al. 2019). We then followed the remainder of the manufacturer’s instructions for DNA isolation and purification.

### Method 2 (M2), DNeasy PowerSoil Isolation Kit supplemented with an enzymatic treatment

We replaced the use of mechanical cell lysis (or bead beating) with a heated, enzymatic treatment step (Yuan et al. 2012), in the DNeasy PowerSoil® DNA Isolation Kit (Qiagen). We added 500 µl of the pooled sample to the kits’ PowerBeads tubes including the tube’s solution (but lacking the beads). We added a lytic cocktail to facilitate cell lysis: 125 µl lysozyme (10 mg/mL, Sigma-Aldrich, St. Louis, MO), 37.5 µl mutanolysin (10 KU/mL Sigma-Aldrich), 7.5 µl lysostaphin (4000 U/mL, Sigma-Aldrich) and 5 µl RNase A (10mg/mL, Sigma-Aldrich). We incubated for 1 hour at 37°C. Then we added 50 µl of proteinase K (20 mg/mL, Sigma-Aldrich) alongside kit solution C1, and followed by incubation for 30 min at 56°C. After centrifugation of the tubes at 10,000 x g for 30 seconds at room temperature (step 6), we continued the rest of the isolation protocol as described by the manufacturer.

### Method 3 (M3), DNeasy UltraClean Microbial Kit

We extracted genomic DNA using the DNeasy UltraClean Microbial Kit (Qiagen) and replaced its bead beating step with the manufacturer’s alternative lysis procedure to reduce shearing of the DNA. We added 500 µl of pooled sample to 300 µl of PowerBead solution. After the addition of 50 µl of Solution SL, we incubated the sample for 10 min at 65°C. After centrifuging the tubes at 10,000 x g for 30 seconds at room temperature (step 5), we continued the rest of the isolation protocol as described by the manufacturer.

### Method 4 (M4), Qiagen Genomic Tip 20/G supplemented with an enzymatic treatment

We extracted DNA using the Qiagen Genomic Tip 20/G (Qiagen) and followed the manufacturer’s protocol “Preparation Gram-negative and some Gram-positive Bacterial Samples”. We centrifuged 500 µl of pooled sample for 10 min at 10,000 x g and resuspended the pellet in Buffer B1 supplemented with 20 µl DNAse-free RNAse (10 mg/mL, Sigma-Aldrich), 45 µl Proteinase K (20 mg/mL, Sigma-Aldrich) and 20 µl lysozyme (100 mg/mL, Sigma-Aldrich), as outlined. We modified the lytic cocktail to include 9 µl lysostaphin (4000 U/mL, Sigma-Aldrich) and 45 µl mutanolysin (10 KU/mL Sigma-Aldrich) in order to improve the lysis potential in gram-positive bacteria. After incubation for 2 hours at 37°C, we added Buffer B2 and extended the incubation at 50°C to 90 min. As the lysate had not cleared after this initial period, we extended the incubation for an additional two hours. We removed any remaining particulate matter by centrifugation at 5,000 x g for 10 min as recommended by the manufacturer. Finally, we used the columns in the kit according to the described isolation protocol.

### Method 5 (M5), Phenol/Chloroform Extraction

We extracted DNA using a phenol/chloroform extraction protocol modified from Chapter 6, protocol 1 of Sambrook and Russell (Sambrook, Jain). For SDS cell lysis, we added 500 µl of pooled sample to 10 mL of TLB (10 mM Tris-HCL pH 8.0, 25 mM EDTA pH 8.0, 100 mM NaCl, 0.5% (w/v) SDS, 20 µg/mL RNase A) and vortexed at full speed for 5 sec followed by an incubation at 37°C for one hour. Proteinase K (Qiagen) was added to a final concentration of 200 µg/mL and we mixed the sample by slow inversion three times, followed by two hours at 50°C with gentle mixing every 30 min. We purified the lysate with 10 mL buffer saturated phenol using phase-lock gel falcon tubes, followed by phenol:chloroform-isoamyl alcohol (1:1). We precipitated the DNA by adding 4 mL of 5 M ammonium acetate and 30 mL ice-cold ethanol. We recovered DNA by one of the following methods: a glass hook followed by washing twice in 70% ethanol (if a DNA mass was visible) or centrifugation at 4500 x g for 10 min followed by washing twice in 70% ethanol (if no DNA mass was visible). After spinning down at 10,000 x g, we removed ethanol by drying at ambient temperature for 10 min. We added 100 µl EB (elution buffer, 10 mM Tris-HCL, pH 8.5) to the DNA and left at 4°C overnight to resuspend the pellet.

### Method 6 (M6), Agarose Plug Extraction

From the 500-ul pooled sample aliquot, we pelleted the cells by centrifugation at 10,000 x g for 10 min and removed the supernatant. We performed DNA extraction on the pellet using a pulsed-field gel electrophoresis (PFGE)-based agarose encasement extraction protocol (Matushek, Bonten, and Hayden 1996). The pelleted cells were resuspended in a 300 µl 2X lysis solution (12.5 mM Tris-HCl pH 7.6, 2 M NaCl, 20 mM EDTA pH 8.0, 1% (w/v) Brij 58, 1% (w/v) deoxycholate, 1% (w/v) sodium lauroyl sarcosine) to which we added lysozyme,1 mg/mL and DNAse-free RNAse, 30 µg/mL on the day of the experiment. We combined the entire suspension with 300 µL of molten 1.6% low-melting point (LMP) agarose (TopVision, Thermo Fisher Scientific, Waltham, MA) which we pipetted into a plug mold and allowed to solidify. We placed each plug into 3 mL of 1X lysis solution containing lysozyme, 0.5 mg/mL; DNAse-free RNAse,100 µg/mL; lysostaphin, 50 U/mL; and mutanolysin, 0.3 KU/mL. We added the enzymes on the day of the experiment and incubated the plugs overnight at 37°C with gentle shaking. After incubation, we replaced the lysis solution sequentially with ESP (10 mM Tris HCl pH 7.6, 1 mM EDTA) to which we added proteinase K (Sigma) at a final concentration of 100 µg/ml and 1% sodium dodecyl sulfate at 50°C for one hour; and then two rounds of washing in sterile, dilute TE (10 mM Tris HCl pH 7.6, 0.1 mM EDTA) first at 50°C for one hour and then at 35°C for 30 min with gentle shaking. We transferred the plug to 5 ml of fresh dilute TE in a clean tube for storage until we performed a ß-agarase I digestion of the agarose and isopropanol DNA precipitation according to the manufacturer’s instructions (New England Biolabs, Ipswich, MA). We resuspended the precipitated DNA in 100 µl EB (10mM Tris-HCL, pH 8.5).

### Determination of DNA yield, purity metrics and fragment size distribution

For each extraction, we quantified the DNA yield on a Qubit® 1.0 Fluorometer (Thermo Scientific), using the dsDNA HS (high sensitivity, 0.2 to 100 ng) Assay kit according to the manufacturer’s protocols; a sample volume of 1 μl was added to 199 μl of a Qubit working solution. We assessed the purity of the extracted nucleic acids with the A260/280 and A260/230 absorbance ratios obtained using a NanoDrop spectrophotometer (NanoDrop Technologies, Wilmington, DE, USA). We assessed the DNA fragment size distribution by electrophoresis (90V for 1.5 h) of genomic DNA (22 ng or 44 ng, as available) on a 0.8% (wt/vol) agarose gel followed by staining with ethidium bromide and UV light visualization. We used λ-HindIII DNA size standards to estimate fragment sizes.

### MinION Library Preparation and Sequencing

Upon receipt, and again immediately prior to sequencing, we measured the flowcell pore counts using the Platform QC script (MinKNOW). We stored the flowcells in their original packaging, which we resealed with parafilm and tape, at 4°C until use. We omitted size selection and the optional shearing step to allow us to evaluate the full distribution of fragment sizes produced by each method. We performed the library preparation using the Ligation Sequencing Kit (SQK-LSK108) and Native Barcoding Kit (EXP-NBD103) for genomic DNA, according to the standard 1D Native barcoding protocol provided by the manufacturer (Oxford Nanopore) (version: NBE_9006_v103_rev2_21DEC16) unless indicated. We carried out all reactions at room temperature. We followed the input DNA mass recommendation of 1.0 µg gDNA and performed DNA repair (NEBNext FFPE DNA Repair Mix, NEB M6630) to maximize read length. For the 1.0x AMPure clean-up step, we used gentle rotation and an extended elution time (minimum 20 min) to assist in the release of long DNA fragments from the beads. We quantified 1 µl aliquots by fluorometry (Qubit) after each phase of the library preparation (*i.e.,* damage repair, end prep, barcoding, pooling and adaptor ligation) to quantify DNA recovery, and identified substantial DNA loss after each step (Table. Quality metrics).

We used two sequencing runs (3 samples multiplexed on each flowcell) for the sequencing. We loaded the completed libraries onto R9.4 flowcells as per instructions from ONT and scheduled sequencing runs for 48 hours. Sequencing continued until time expired or until pore exhaustion (defined by <10 functional pores).

### BluePippin Size Selection

We performed size selection by the BluePippin (Sage Science) system with 0.75% dye-free agarose cassettes and marker S1. We selected fragments > 6 kB in high-pass collection mode (an approach enabling the collection of DNA fragments above a user-defined size). Our 3.3 µg sample DNA input was less than the recommended 5 µg; however, this input was maximized given sample DNA concentrations and loading volume constraints.

### Long-read Sequence analysis

We uploaded the raw MinION FAST5 files produced with the MinKNOW software (versions 1.15.4 through 3.1.19) to our cluster to perform the base-calling and demultiplexing with guppy v2.3.1. FastQ files were generated only for reads meeting a minimum quality threshold (quality score of 7). We used minimap2 v2.14 (Li 2018) and samtools v1.9 (Li et al. 2009) to remove sequences that mapped to the human genome build 38 (GRCh38) from NCBI and estimate the amount of host contamination. We used anvi’o v6.2 and the contig snakemake workflow to compute the sequence metrics (Köster and Rahmann 2012). Briefly, the workflow created a contigs database with ‘anvi-gen-contigs-database’, which used Prodigal v2.6.3 (Hyatt et al. 2010) with the metagenome mode to identify open reading frames. It used ‘anvi-run-hmm’ to detect the single-copy core genes from bacteria (n=71, modified from (Lee 2019)), archaea (n=76, (Lee 2019)), eukarya (n=83, http://merenlab.org/delmont-euk-scgs), and ribosomal RNAs (n=12, modified from https://github.com/tseemann/barrnap). We used ‘anvi-display-contigs-stats’ to get the number of sequences, total length, N50, longest sequence, number of genes, number of single-copy core genes and ribosomal genes. We used ‘anvi-get-sequences-for-hmm-hits’ to recover 16S rRNA genes for each condition and used the Human Oral Microbiome Database (HOMD) online blast tool for taxonomic assignment. We assembled the long reads with Flye and the metagenomic option (Kolmogorov et al. 2019), which takes into account the uneven coverage nature of metagenomes. We created anvi’o contigs databases, as described above, with Flye’s contigs to summarize the assembly metrics.

### 16S rRNA gene amplicon DNA extraction, library preparation, sequencing and analysis

We performed sample DNA extraction using the DNeasy Powersoil kit (Qiagen) following the manufacturer’s protocol. We amplified the V4-V5 hypervariable regions of the bacterial SSU rRNA gene using degenerate primers. 518F (CCAGCAGCYGCGGTAAN) and 926R (CCGTCAATTCNTTTRAGT, CCGTCAATTTCTTTGAGT, and CCGTCTATTCCTTTGANT). Amplification was done with fusion primers containing the 16S-only sequences fused to Illumina adapters. The forward primers included a 5 nt multiplexing barcode and the reverse a 6 nt index. We generated PCR amplicons in triplicate 33 µl reaction volumes with an amplification cocktail containing 0.67 U SuperFi Taq Polymerase (Invitrogen, Carlsbad, CA), 1X enzyme buffer (includes MgCl2), 200 µM dNTP mix (ThermoFisher), and 0.3 µM of each primer. We added approximately 10-25 ng template DNA to each PCR and ran a no-template control for each primer pair. Amplification conditions were: initial 94C, 3 minute denaturation step; 30 cycles of 94C for 30s, 57C for 45s, and 72C for 60s; final 2 minute extension at 72C. The triplicate PCR reactions were pooled after amplification, visualized with the negative controls on a Caliper LabChipGX, and purified using Ampure followed by PicoGreen quantitation and Ampure size selection. Libraries were sequenced on an Illumina Miseq 250-cycle paired-end run. We used illumina-utils v2.7 (Eren et al. 2013) for the quality filtering, following (Minoche, Dohm, and Himmelbauer 2011) recommendations, and the merging of the paired-end reads. We used Vsearch to remove chimeric sequences (Rognes et al. 2016) and Minimum Entropy Decomposition (MED) (Eren et al. 2015) to cluster the merged reads into oligotypes. We assigned taxonomy using DADA2 (Callahan et al. 2016) and the Silva v132 non-redundant database (Quast et al. 2013).

### Short-read metagenomic library preparation, sequencing and analysis

Sample DNA concentrations, determined by PicoGreen assay, were 67 ng/ul (TD) and 0.75 ng/ul (M1_2). We used 100 ng and 28 ng, respectively, for library construction. DNA was sheared to ∼400 bp using the Covaris S2 acoustic platform and libraries were constructed using the Nugen Ovation Ultralow kit. Each required an amplification step: 7 cycles (TD) or 11 cycles (M1_2). The products were visualized on an Agilent Tapestation 4200 and size-selected to an average of 482 bp using BluePippin (Sage Biosciences). The final library pool was quantified with the Kapa Biosystems qPCR protocol and sequenced on the Illumina NextSeq500 in a 2 × 150 paired-end sequencing run using dedicated read indexing. We used anvi’o v6.2 and the metagenomics snakemake workflow for the assembly and analysis of the short-reads. Briefly, the workflow uses illumina-utils for quality filtering followed by a metagenomics assembly with IDBA-UD v1.1.3 (Peng et al. 2012) and generates contigs databases as described in the long-read sequence analysis section. The workflow also uses Bowtie2 v2.3.5.1 (Langmead and Salzberg 2012) to map short-read on the assembled contigs and samtools (Li et al. 2009) to sort, index and convert sam files into bam files used by anvi’o to generate profiles databases. To compute taxonomic profiles of metagenomes we used the anvi’o program ‘anvi-estimate-scg-taxonomy’, which aligns ribosomal proteins found in a metagenomic assembly to those that are found in reference genomes from the Genome Taxonomy Database (GTDB) (Parks et al. 2020).

### Visualization

We generated the figures in R with ggplot2 (Wickham 2009) and modified them with Inkscape.

### Data availability

Raw sequences for long- and short-read sequencing data for shotgun metagenomes and 16S rRNA gene amplicons are available under the BioProject PRJNA703035. We also made available FASTA files and anvi’o contigs databases for (1) assembled long-read sequences at doi:10.6084/m9.figshare.14141228, (2) unassembled long-read sequences at doi:10.6084/m9.figshare.14141414, and (3) assembled shotgun metagenomes at doi:10.6084/m9.figshare.14141819. Supplementary Tables and Figures are also available via doi:10.6084/m9.figshare.14141918.

## Acknowledgements

We thank Nicole Robichaud for technical help with short-read sequencing. This research was supported by the Gastro-Intestinal Research Foundation, the Simons Foundation (#687269, AME). In addition, a Helmsley Foundation grant to LB, BJ, and AME supported FT, and an NIH NIDDK grant (RC2 DK122394) supported KL and AME.

## Author contributions

FT, KL, and AME conceived the study. KL developed and implemented long-read sequencing protocols. FT performed analyses of short- and long-read sequencing data. KL, EF, and AS helped with data analysis. HGM performed short-read sequencing. HGM, LB, BJ, and AME supervised research. FT, KL, and AME wrote the paper with critical input from all authors.

---

**Figure S1.**
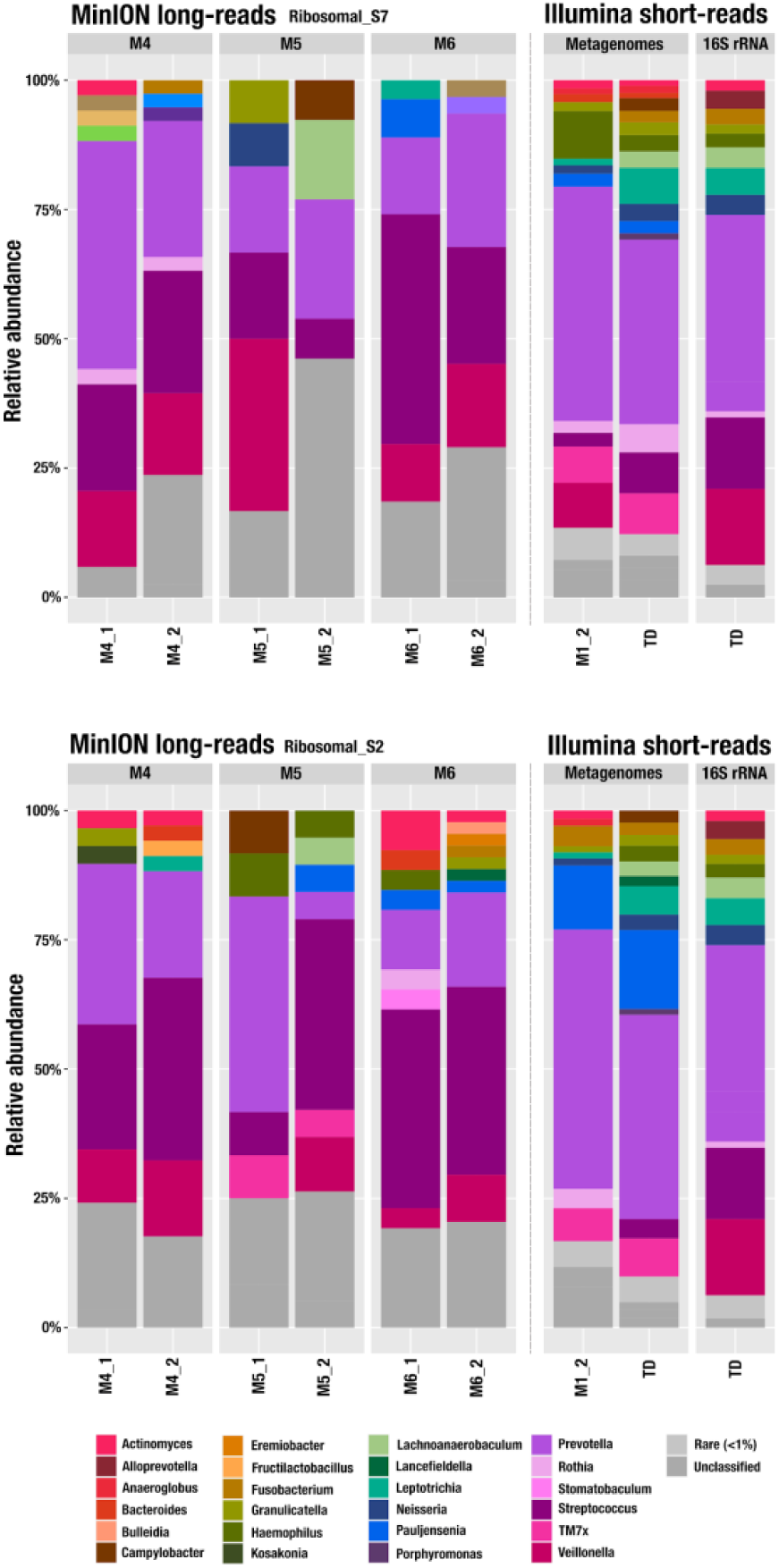
Taxonomic relative abundance at the genus level with ribosomal protein S7 and S2. We used the GTDB database to assign taxonomy to the two most abundant ribosomal proteins (S7 and S2) identified in long-read metagenomes using HMMs. For the short-read metagenomes, we used the taxonomy of the ribosomal gene S7 with the Genome Taxonomy Database (GTDB). We performed a 16S rRNA amplicon sequencing on an additional sample (Amplicon) and used MED with Silva (v132) assignation at the genus level. Genera representing less than 1% of a sample were pooled as rare (light grey).

**Table S1**. Read counts and cumulative length. (a) Number of sequences (and the cumulative length) of the MinION outputs, before and after quality filtering. We used a qscore of 7 as the quality filtering threshold. (b) Microbial read size distribution metrics. All percentages are relative to the total amount of reads (or total length) of the sequencing output, before filtering out human contamination, but after quality filtering (criteria: minimum Q-score of 7). (c) Impact of size selection on read-size distribution.

**Table S2**. Oligotypes identified by Minimum Entropy Decomposition (MED). (a) Matrix count, (b) representative sequences and (c) associated Silva taxonomy.

**Table S3**. Distribution of ribosomal proteins across assembled long-reads across extraction strategies.

